# Lipopolysaccharide synthesis and traffic in the envelope of the pathogen *Brucella abortus*

**DOI:** 10.1101/2022.05.19.492625

**Authors:** Caroline Servais, Victoria Vassen, Audrey Verhaeghe, Nina Küster, Elodie Carlier, Léa Phégnon, Aurélie Mayard, Xavier De Bolle

**Author notes:** Corresponding author, Address: 61 Rue de Bruxelles, 5000 Namur, Belgium.

## Abstract

Lipopolysaccharide is essential for most Gram-negative bacteria as it is a main component of the outer membrane. In the pathogen *Brucella abortus*, smooth lipopolysaccharide containing the O-antigen is required for virulence. Being part of the Rhizobiales, *Brucella* spp. display unipolar growth and lipopolysaccharide was shown to be incorporated at the active growth sites, *i*.*e*. the new pole and the division site. By localizing proteins involved in the lipopolysaccharide transport across the cell envelope, from the inner to the outer membrane, we show that the lipopolysaccharide incorporation sites are determined by the inner membrane complex of the lipopolysaccharide transport system. Moreover, we identify the main O-antigen ligase of *Brucella spp* involved in smooth lipopolysaccharide synthesis. Altogether, our data highlight a new layer of spatiotemporal organization of the lipopolysaccharide biosynthesis pathway and identify a new class of bifunctional O-antigen ligases.

## Main

The Gram-negative cell envelope is composed of an inner membrane (IM), a periplasmic space hosting peptidoglycan (PG), and an outer membrane (OM) (Beveridge, 1999). The OM constitutes the interface with the external environment. It is an asymmetric bilayer mainly composed of phospholipids in the inner leaflet and lipopolysaccharide (LPS) in the outer leaflet (Silhavy *et al*., 2010; Bertani *et al*., 2018). LPS is essential for OM integrity in most Gram-negative bacteria (Zhang *et al*., 2013). This amphiphilic molecule is composed of lipid A, a hydrophobic anchor made of acyl chains, a core composed of sugars, and a long polysaccharidic chain, called the O-antigen. Depending on the presence or the absence of the O-antigen, the LPS is referred to as smooth (S-LPS) or rough (R-LPS). Most rod-shaped bacteria grow and elongate their cell wall in a dispersed manner as the Gram-negative model organism *Escherichia coli* (Burman *et al*., 1983; Woldringh *et al*., 1987). In contrast, bacteria from the Rhizobiales order belonging to the class of alpha-proteobacteria, such as *Brucella abortus, Agrobacterium tumefaciens*, and *Sinorhizobium meliloti*, grow unipolarly (Brown *et al*., 2012). New envelope material is incorporated at one pole of the cell (namely the new pole), resulting in a daughter cell exclusively composed of new envelope material (Brown *et al*., 2012). Several proteins were shown to polarly localize at the new or old pole, in agreement with the association of distinct functions at the poles. Among others, two Rhizobiales <u>g</u>rowth and <u>s</u>eptation (Rgs) proteins, RgsS and RgsE were recently localized in *S. meliloti*. RgsS, a FtsN-like protein, was shown to be localized at the growth sites, while RgsE, a homolog of the Growth Pole Ring protein (GPR) of *A. tumefaciens* (Zupan *et al*., 2019), was shown to be localized exclusively at the new pole (Krol *et al*., 2020; Krol *et al*., 2021). Both proteins are thought to be important to maintain proper polar growth.

Brucellosis is a worldwide zoonosis mainly caused by *B. abortus, Brucella melitensis* and *Brucella suis*. These three species are very close at the phylogenetic level but differ by their host specificity (Moreno *et al*., 2006). However, they display S-LPS at the surface linked to a virulent phenotype. In *B. abortus*, PG, two OM proteins (OMP) and LPS were shown to be incorporated at the growth sites, *i. e*. the new pole and the division site (Vassen *et al*., 2019). Moreover, *B. abortus* also displays a mixture of R-LPS and S-LPS on the surface of the smooth wild type (Vassen *et al*., 2019). Interestingly, *Brucella* LPS is different from the classical enterobacterial LPS (Lapaque *et al*., 2005), as for instance, *Brucella* lipid A has one longer acyl chain and its core has a branched structure (Lapaque *et al*., 2005; Conde-Alvarez *et al*., 2012). During LPS biosynthesis, lipid A and the core are assembled at the inner leaflet of the IM to form the R-LPS that is subsequently flipped to the outer leaflet of the IM by the essential ABC transporter MsbA (Bonifer *et al*., 2021). The O-antigen, a long unbranched homopolymer of 4,6-dideoxy-4-formamido-α-D-mannopyranosyl (*N*-formyl-perosamine) found in most *Brucella* strains (Caroff *et al*., 1984; Wattam *et al*., 2012), is the most variable part of the LPS. Its length highly varies within a *Brucella* population and even on individual bacterial cell (Dubray *et al*., 1987; Bowden *et al*., 1995). During its biosynthesis, the O-antigen is linked to a lipid carrier, a 55-carbon isoprenoid, the undecaprenyl phosphate (Und-P) (Whitfield *et al*., 2014), at the inner leaflet of the IM and is subsequently flipped by a different ABC transporter complex, Wzm/Wzt (Godfroid *et al*., 2000; Bi *et al*., 2018). At the periplasmic side of the IM, an unknown O-antigen ligase links the O-antigen to a fraction of R-LPS to generate S-LPS. R-LPS and S-LPS are then translocated to the OM via the <u>LP</u>S transport machinery (Lpt). The Lpt complex forms a bridge of seven proteins (LptAB2CDEFG), spanning from the IM to the OM (Sperandeo *et al*., 2017). It has been mainly studied in *E. coli* and all of the proteins constituting the machinery are conserved and essential in *B. abortus* (Sternon *et al*., 2018). Until now, the Lpt proteins have never been localized in unipolarly growing bacteria and the O-antigen ligase remains unknown in *Brucella spp*.

Here, we investigate the localization of proteins involved in unipolar growth in *B. abortus*. We first demonstrate that the polar anchoring of two Rgs proteins is conserved between *S. meliloti* and *B. abortus*. In adddition, we localize several components of the Lpt machinery in the IM and the OM, as well as the lipidA-core flippase, MsbA. We identify WadA as a bifunctional enzyme and the main O-antigen ligase in *Brucella* spp. WadA thereby displays a cytoplasmic glycosyltransferase activity adding the last core sugar necessary for the O-antigen ligation, in addition to the O-antigen ligase activity that is proposed to take place in the periplasmic space. Altogether, we propose a model for LPS biosynthesis that involves LPS trafficking from the old pole in the IM to the new pole in the OM of *B. abortus*.

### Polar localization of the essential RgsS and RgsE proteins is conserved in *B. abortus*

Before localizing enzymes and transporters involved in LPS insertion in the OM, we wondered if proteins identified at the growth pole in *S. meliloti*, RgsS and RgsE, were conserved and localized at similar sites in *B. abortus*. RgsS and RgsE homologs in *B. abortus* were genetically fused to mNeonGreen (mNG) and expressed from their normal chromosomal locus. Since both genes are essential in *B. abortus* (Sternon *et al*., 2018), viable mutants displaying mainly a normal morphology suggests that these fusions are functional. RgsS-mNG and RgsE-mNG were analyzed in a strain producing PdhS-mCherry as old pole marker (**Fig. 1**) (Hallez *et al*., 2007). Demograph analysis revealed that RgsS was localized at the new pole in smaller bacteria and at the mid cell in predivisional bacteria, while RgsE was exclusively localized at the new pole (**Fig. 1**). These localization patterns are comparable to *S. meliloti*, suggesting that at least a fraction of Rhizobiales share similar localization of proteins involved in unipolar growth.

**Figure 1.**
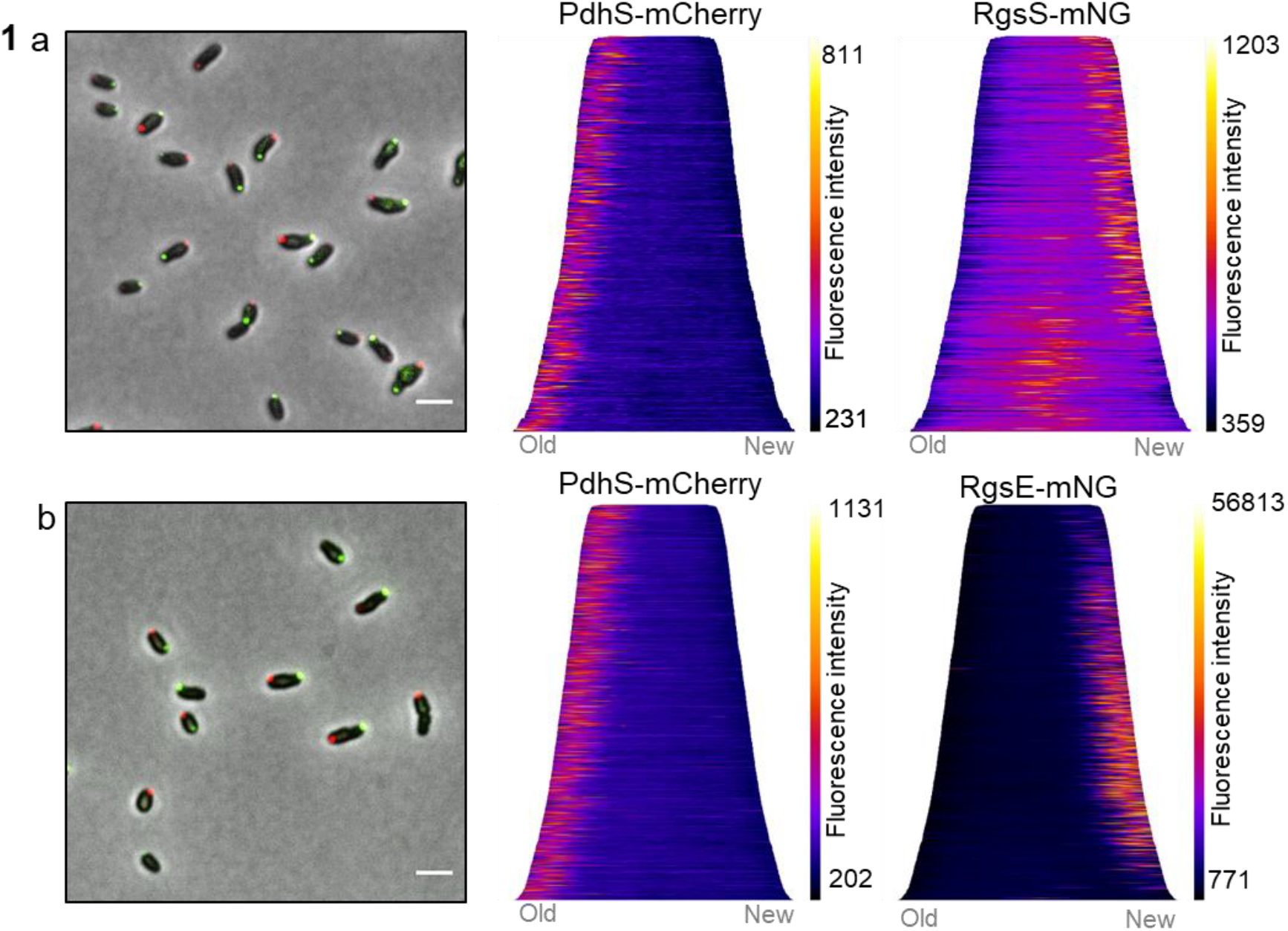
Polar localization of Rhizobiales growth and septation (Rgs) proteins is conserved in *B. abortus*. Representative merged picture with phase and fluorescent channels of the strain co-expressing PdhS-mCherry (red) and (a) RgsS-mNG (green) or (b) RgsE-mNG. Scale bar is 2μm. The right panels show demographic representations of PdhS-mCherry and of (a) RgsS-mNG or (b) RgsE-mNG localization. Cells are aligned based on their length (smallest on top) and pole age using PdhS-mCherry as an old pole marker (old pole on the left). The graphs correspond to one representative experiment (n=449 bacteria for RgsS-mNG and n=1482 bacteria for RgsE-mNG). Fluorescence intensity is represented as a heatmap, the minimum and maximum value represented on the scale were automatically selected to provide the best signal to background by MicrobeJ.

### The OM component of the Lpt pathway, LptD, is dispersed on *B. abortus* surface

Given that LPS is integrated unipolarly in the OM (Vassen *et al*., 2019), we wonder whether the LPS translocation pathway was also localized at the growing cell pole. Together with LptE, LptD forms the OM part of the Lpt pathway, allowing the last step of the LPS transport to the OM (Wu *et al*., 2006; Botte *et al*., 2022).In order to localize this large beta barrel in *B. abortus*, a 3Flag tag was incorporated into an unconserved loop of LptD (Fig. S1), predicted to be surface exposed (Fig. 2a). The *lptD* gene was replaced by the engineered *3Flag::lptD* fusion and the resulting strain was viable and the fusion protein was detectable (Fig. 2b). Since *lptD* is essential, viability and normal morphology of the engineered strain suggest that 3Flag::LptD is functional. The fusion protein was localized by scanning electron microscopy using an anti-3Flag monoclonal antibody (mAb) and a secondary antibody conjugated to gold particles. The fusion could only be detected on the surface of the rough mutant strain (disrupted *gmd*) and not in the wild type (WT) strain, presumably because the long O-antigen impaired the detection of the surface exposed 3Flag tag. The *gmd* mutant with endogenous *lptD* gene was used as negative control to eliminate the gold particles background, *i*.*e*. cells with ≤ 4 gold particles (Fig. S2a & S2b). The distribution frequency of the gold particles showed that LptD is almost evenly located on the bacterial surface, with a slightly lower frequency near the constriction site (Fig. S2c). In contrast to the polar insertion of LPS, the apparent homogenous LptD localization could be explained by the limited mobility of OMP as previously shown (Vassen *et al*., 2019; Benn *et al*., 2021). We therefore propose that LptD is only active at the growth sites, receiving new LPS molecules by the Lpt proteins located in the IM.

**Figure 2.**
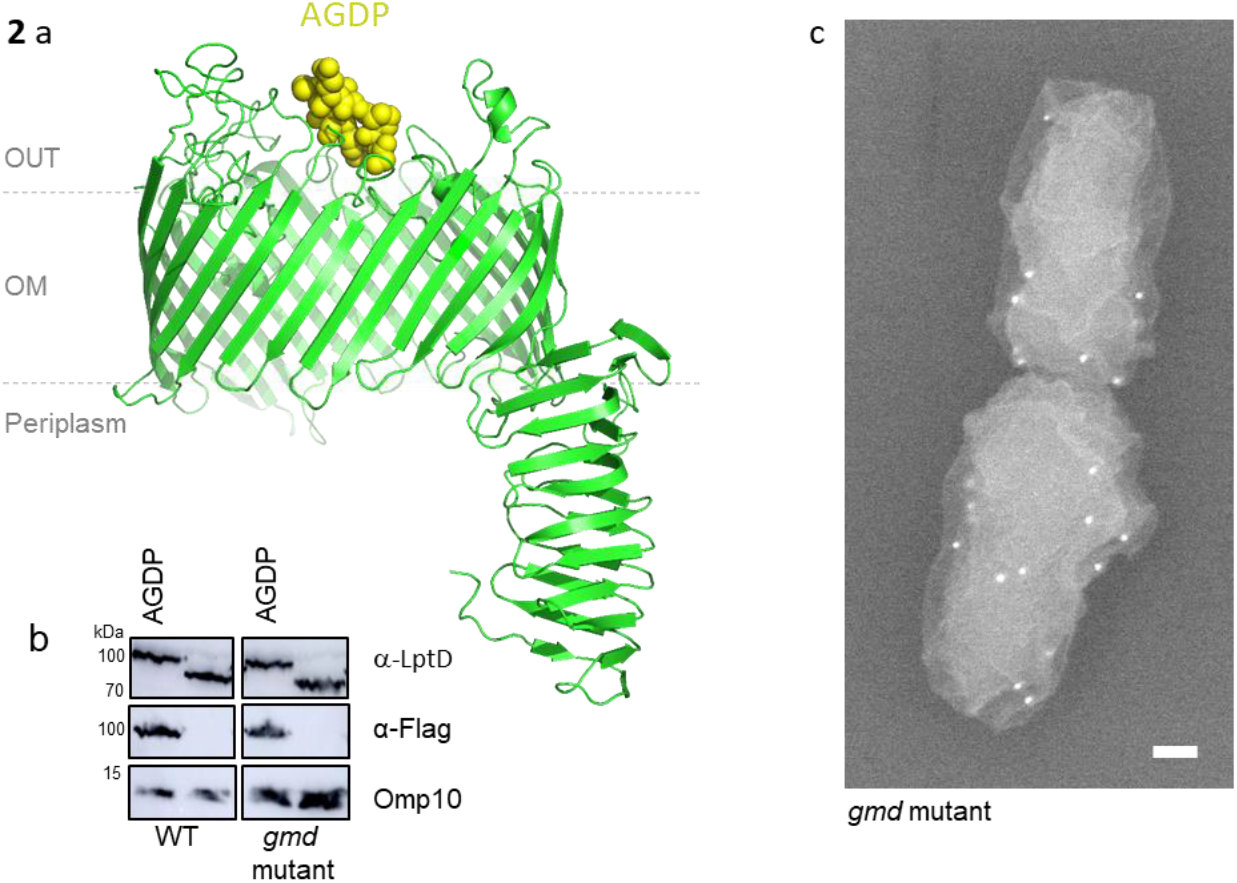
3Flag::LptD is found dispersed on the *B. abortus* cell surface. **a**, Three-dimensional model of LptD highlighting the ADGP loop (yellow spheres) in which the 3Flag was inserted. This sequence was among the most variable in the highly conserved LptD sequence (Fig. S1). The 3D structure of LptD was predicted using Swiss model server (https://swissmodel.expasy.org) (Waterhouse *et al*., 2018) and displayed with PyMol v.2.0 (Schrodinger, 2015) **b**,The presence of the 3Flag in LptD was assessed by western blot (WB) in the WT and in the *gmd* mutant with polyclonal antibodies directed against LptD and mAb recognizing the 3Flag. Omp10 was used as a loading control. **c**, Representative scanning electron microscopy picture of *gmd* mutant with 3Flag::lptD fusion. 3Flag-LptD was detected using mAb against 3Flag, followed by secondary antibody coupled with gold particles, each white focus corresponds to a single gold particle with the expected size (18 nm). Scale bar represents 100 nm.

### The IM components of the Lpt complex are localized at the growth sites

The IM components of the Lpt transport machinery include LptB, LptC, LptF and LptG (Sperandeo *et al*., 2017), which all have a distinct and essential homolog in *B. abortus* (Sternon *et al*., 2018). The localization of LptC was achieved by fusing mNG at the N-terminus (*mNG-lptC*), *i*.*e*. the cytoplasmic part of the protein, by allelic replacement at the chromosomal locus. The *mNG-lptC* strain was viable, grew similarly to the WT in rich medium and still displayed unipolar growth indicating a functional fusion (Fig. S3a & b). The old pole marker PdhS-mCherry was used to determine the old pole. LptC was mainly found at the new pole in non-divisional bacteria, or at the mid cell in pre-divisional cells (Fig. 3a), similar to the LPS incorporation profile. Demographic analysis confirmed this localization pattern (Fig. 3a). Since LptC forms a stable complex with LptB_2_FG in other bacteria (Narita *et al*., 2009), we also investigated the localization of mNG-LptF to confirm that another component of the IM complex of the Lpt system is co-localizing with LPS insertion sites. The strain *mNG-lptF* was constructed by allelic replacement and showed unipolar growth (Fig. S3). The mNG-LptF localization pattern was comparable to mNG-LptC (Fig. 3b), further supporting the hypothesis that LptD localized at the OM is provided with LPS from Lpt complex localized in the IM.

**Figure 3.**
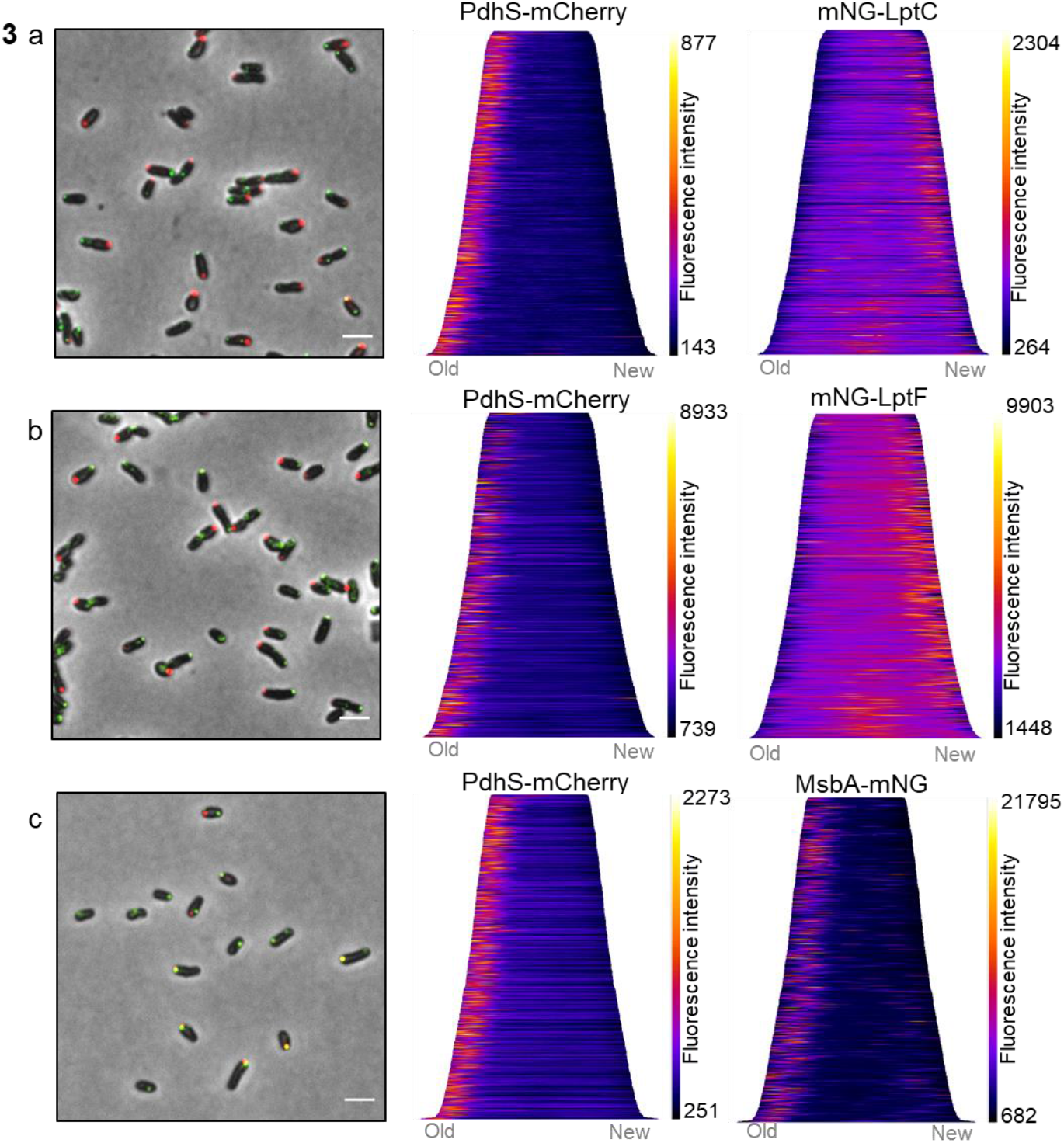
LptC, LptF and MsbA localization. Representative merged microscopy pictures and demograph analyses of (a) *mNG-lptC* (n=723 bacteria), (b) *mNG-lptF* (n=1398 bacteria), and (c) *msbA-mNG* (n=693 bacteria) co-expressing the old pole marker PdhS-mCherry. The mNG-LptC (a) and mNG-LptF (b) are detected at the growth sites while MsbA-mNG is localized at the old pole (c). Scale bars represent 2 μm. The right panels show demographic representation. The cells were sorted according to their size (from the smallest on top to the largest at the bottom) and oriented according to their cell poles using the old pole marker PdhS-mCherry. Demographs are representative images. Fluorescence intensities are represented as heatmap with minimum and maximum values represented on the vertical-scale, and were automatically selected to provide best signal to background ratio using MicrobeJ image analysis.

### The essential flippase MsbA is mainly found at the old pole

Since the IM Lpt proteins are localized at the growing zones, it was tempting to hypothethise that the molecular actors involved in the LPS biosynthesis upstream of the Lpt, such as MsbA, would be in a close vicinity to facilitate LPS translocation through the Lpt system. Therefore, a *msbA-mNG* fusion was constructed and integrated at the chromosomal locus to replace the endogenous essential *msbA* gene. As previously, the genetically modified strain was viable, displayed a normal growth in rich medium as well as unipolar growth (Fig. S3), suggesting that MsbA-mNG is functional. MsbA-mNG localization was analyzed in a *pdhS-mCherry* background. Surprisingly, MsbA was found at the old pole in 97% of the bacteria (Fig. 3c and Fig. S4), which was in total contradiction with our expectation. Manual counting highlighted that 52% of the bacteria only presented one focus at the old pole, while 47% of them presented at least one additional clearly distinct focus throughout the whole bacterium in addition to the old pole (Fig. S4).

In a next step, we were wondering if MsbA activity was coupled to its old pole localization. To address this question, we generated a strain in which *msbA* was duplicated. The strain contains an intact copy of *msbA* together with a *msbA-mNG* fusion (pSK_*msbA-mNG*) and the PopZ-mCherry new pole marker (Deghelt *et al*., 2014). This strain displayed normal growth and MsbA localization pattern comparable to the strain with a single *msbA-mNG* gene (Fig. 4a). Furthermore, we constructed a strain in which the additional *msbA-mNG* gene was replaced with a *msbA-mNG* allele in which the glutamate 491 codon was replaced by an alanine (pSK_*msbA*_E491A_-*mNG*). As already described for other ABC transporters, mutating the corresponding glutamate on the Walker B Motif impairs ATP hydrolysis, thereby blocking the protein in an inactive ATP-bound conformation (Orelle *et al*., 2003). Interestingly, MsbA_E491A_-mNG localization was dispersed and remaining foci were much less intense (Figure 4b), suggesting that the proper localization of MsbA is coupled to its ATPase activity.

**Figure 4.**
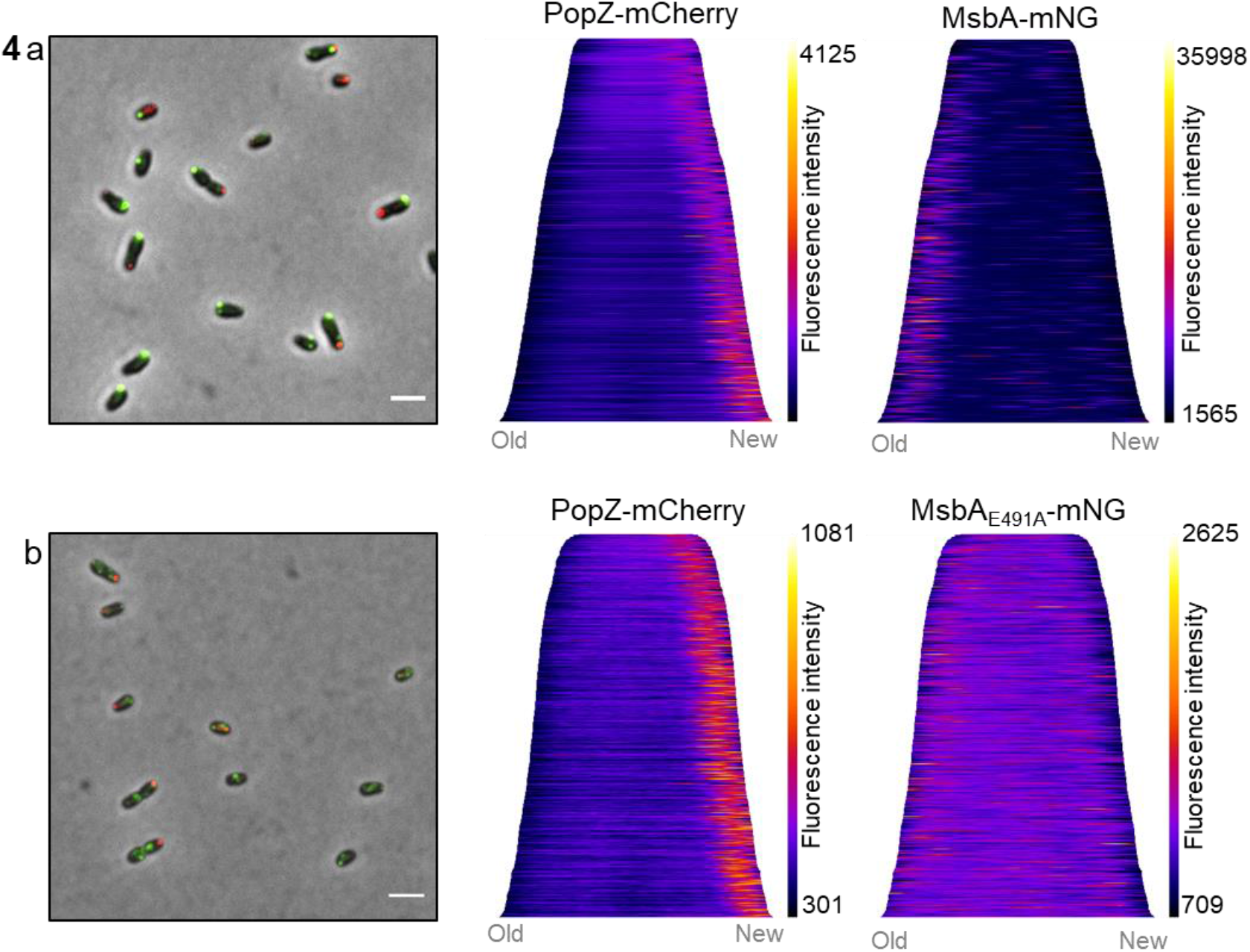
Mutation in the Walker B motif of MsbA leads to diffused localization. Representative merge pictures and demograph analyses for two strains **a**, an additional copy of. MsbA fused to mNG (pSK_*msbA-mNG*) was still mainly localized at the old pole. The two demographs represent PopZ-mCherry (new pole marker) and MsbA-mNG localization (n=1095 bacteria). **b**, an additional copy of MsbA mutated for E491A and fused to the mNG. MsbA_E491A_-mNG (pSK_*msbA*_*E491A*_*-mNG*) showed general less intense and more dispersed signal. Demograph analyses represent PopZ-mCherry and MsbA_E491A_-mNG localization respectively (n=536 bacteria). Fluorescence intensities are represented as heatmaps with automatically selected minimum and maximum values represented on the scale in order to provide the best signal to background ration by MicrobeJ. Scale bars are 2μm.

### WadA is the O-antigen ligase in *Brucella* species

Although the whole LPS translocation pathway has been identified, one important biosynthesis enzyme was still missing. Indeed, the O-antigen ligase, attaching the O-antigen to the lipid A-core using Und-PP-O-antigen as a substrate, remains unknown in *Brucella* spp. We identified two putative candidates, WaaL and WadA, that were predicted to code for O-antigen ligases based on domain composition. WaaL has been identified as the main O-antigen ligase in many bacterial species (Schild *et al*., 2005; Islam *et al*., 2010; Li *et al*., 2017; Perez-Burgos *et al*., 2019). However, deletion of the *waaL* homolog in *B. abortus* resulted in smooth phenotype like the wild type (Fig. S5), suggesting that WaaL is not the main O-antigen ligase under the tested conditions.

Three dimensional structure prediction of *Brucella* WadA using Alpha-Fold (Jumper *et al*., 2021; Varadi *et al*., 2022) highlighted 12 transmembrane domains with periplasmic loops (Fig. S6), similar to the *E. coli* O-antigen ligase topology (Perez *et al*., 2008; Ashraf *et al*., 2022). The *wadA* gene was scored as essential in *B. abortus* 2308 (Sternon *et al*., 2018), while its ortholog was not essential in *B. melitensis* (Gonzalez *et al*., 2008; Salvador-Bescos *et al*., 2018). However, these two *wadA* sequences are identical on a sequence level. In *B. melitensis*, WadA acts as the glycosyltransferase that adds the terminal core sugar (a glucose) needed as attachment structure for the O-antigen (Salvador-Bescos *et al*., 2018). The rough phenotype of *B. melitensis wadA* deletion mutant (Δ*wadA*) was confirmed both by western blot (WB) analysis (Fig 5a) and immunofluorescence microscopy (Fig. S7). WadA contains two domains, a cytoplasmic N-terminal glycosyltransferase (GT) domain and a predicted C-terminal O-antigen ligase (OAg-lig) domain. Deletion of a major part of the largest periplasmic loop of the *OAg-lig* domain (Δ*OAg-lig*) was constructed (see Material and Method). The Δ*OAg-lig* mutant displayed a rough phenotype detected by WB analysis using the core targeting antibody (A68/24D08), allowing the distinction between R-LPS and S-LPS after electrophoresis. The Δ*OAg-lig* mutant could be complemented by adding a copy of the *OAg-lig* domain coding sequence, suggesting that the OAg-lig domain is indeed coding for a functional O-antigen ligase in *B. melitensis*.

**Figure 5.**
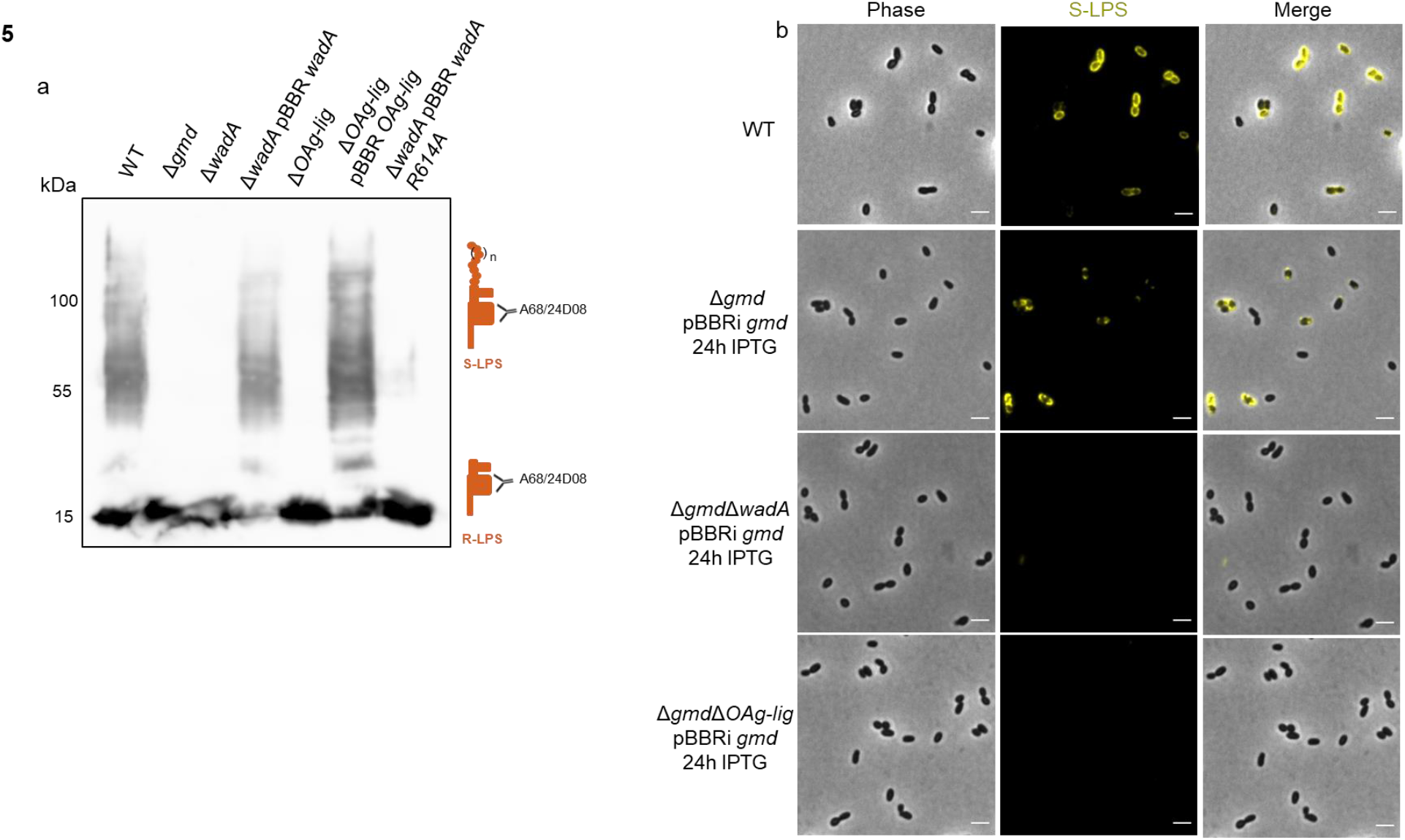
WadA is the main O-antigen ligase in *Brucella* spp. **a**, WB analysis using an antibody against the core of the LPS (A68/24D08) in *B. melitensis* 16M mutants. **b**, Immunofluorescence microscopy labeling the S-LPS (A76/12G12) of the WT *B. abortus* 544, Δ*gmd*Δ*wadA*, and the Δ*gmd*Δ*OAg-lig* expressing *gmd* upon IPTG induction for 24h. Scale bars are 2μm.

To further support the role WadA as the main O-antigen ligase in *Brucella*, we mutated an arginine (Arg-614) conserved in several WadA homologs among the Rhizobiales (Fig. S6). Arg 614 is located at the terminal region of the largest periplasmic loop of the OAg-lig domain (Fig. S6). Indeed, analysis of *E. coli* O-antigen ligase revealed a cluster of positively charged residues in the periplasmic loop that were required for O-antigen ligation (Perez *et al*., 2008; Ruan *et al*., 2012). Complementation of Δ*wadA* with *wadA* R614A lead to very low levels of S-LPS synthesis, while complementation with intact *wadA* restores the WT S-LPS phenotype (Fig. 5a & Fig. S7). A recent study showed that several other residues were involved in O-antigen ligation (Ashraf *et al*., 2022), which could explain why some residual S-LPS can be detected in the presence of the WadA-R614A mutant.

It was previously shown that alteration of the O-antigen synthesis leads to growth defects in *E. coli* (Jorgenson *et al*., 2016). Since the Und-P lipidic carrier binds both O-antigen and PG precursors, alteration of the free Und-P pool could also impair PG synthesis and among others bacterial growth (Jorgenson *et al*., 2016). We therefore hypothesized that the essentiality of *wadA* in *B. abortus* could be due to an indirect effect of Und-P sequestration by the O-antigen. Indeed, *ΔwadA* and *ΔOAg-lig* deletions strains could be obtained in a Δ*gmd* strain unable to synthesize the O-antigen. In order to show that WadA and more particularly its OAg-lig domain is required for the production of S-LPS in *B. abortus*, we introduced the *gmd* gene in the Δ*gmd*Δ*wadA* and Δ*gmd*Δ*OAg-lig* strains, to test the hypothesis that Δ*wadA* and Δ*OAg-lig* mutations would impair S-LPS synthesis even when *gmd* is expressed. A copy of *gmd* was added under the control of the p_*lac*_ promoter in the Δ*gmd*, Δ*gmd*Δ*wadA* and Δ*gmd*Δ*OAg-lig* strains. As expected, immunofluorescence microscopy after induction of *gmd* expression showed no fluorescence signal for S-LPS neither in Δ*gmd*Δ*wadA* nor in Δ*gmd*Δ*OAg-lig* whereas S-LPS labelling could be detected in Δ*gmd* after induction used as positive control (Fig 5b).

During growth, Gram-negative bacteria must expand the different layers of their cell wall, PG and OM, in a temporal and spatial coordinated manner. Similar to other bacteria of the Rhizobiales order, *B. abortus* displays unipolar growth incorporating new envelope components at the new pole in non-divisional bacteria and at the constriction site of divisional bacteria. Previously, LPS was one of the components reported to be unipolarly incorporated at the OM of *B. abortus* (Vassen *et al*., 2019). However, to the best of our knowledge, the essential LPS translocation pathway has never been localized before. Here, we show that although the main OM component of the Lpt pathway, LptD, is found dispersed at the bacterial surface, the IM components LptC and LptF are localized at the growth sites, in accordance with the LPS incorporation sites (Fig. 6). How to explain that strikingly, LptD localization does not align with the LPS insertion sites? When a component like LptD is inserted in the OM, it would remain fixed at its incorporation site in the case of a static OM. This was previously suggested by the analysis of Omp2b porin, Omp25 and LPS (Vassen *et al*., 2019), and by the covalent attachment of several OMPs to the PG (Godessart *et al*., 2021). LptD would only incorporate LPS during growth at the growth sites when it receives substrates from the IM Lpt proteins such as LptC and LptF shown to be localized at the growth sites. How LptC and LptF are targeted to the growth sites and how LptD can switch from an inactive to an active state remains to be discovered. It is not excluded that LptD proteins located outside the canonical growth areas switch from an inactive to an active form, for example in strains that produce blebs and thus need to compensate the loss of OM fragments everywhere on their surface (Godessart *et al*., 2021).

**Figure 6.**
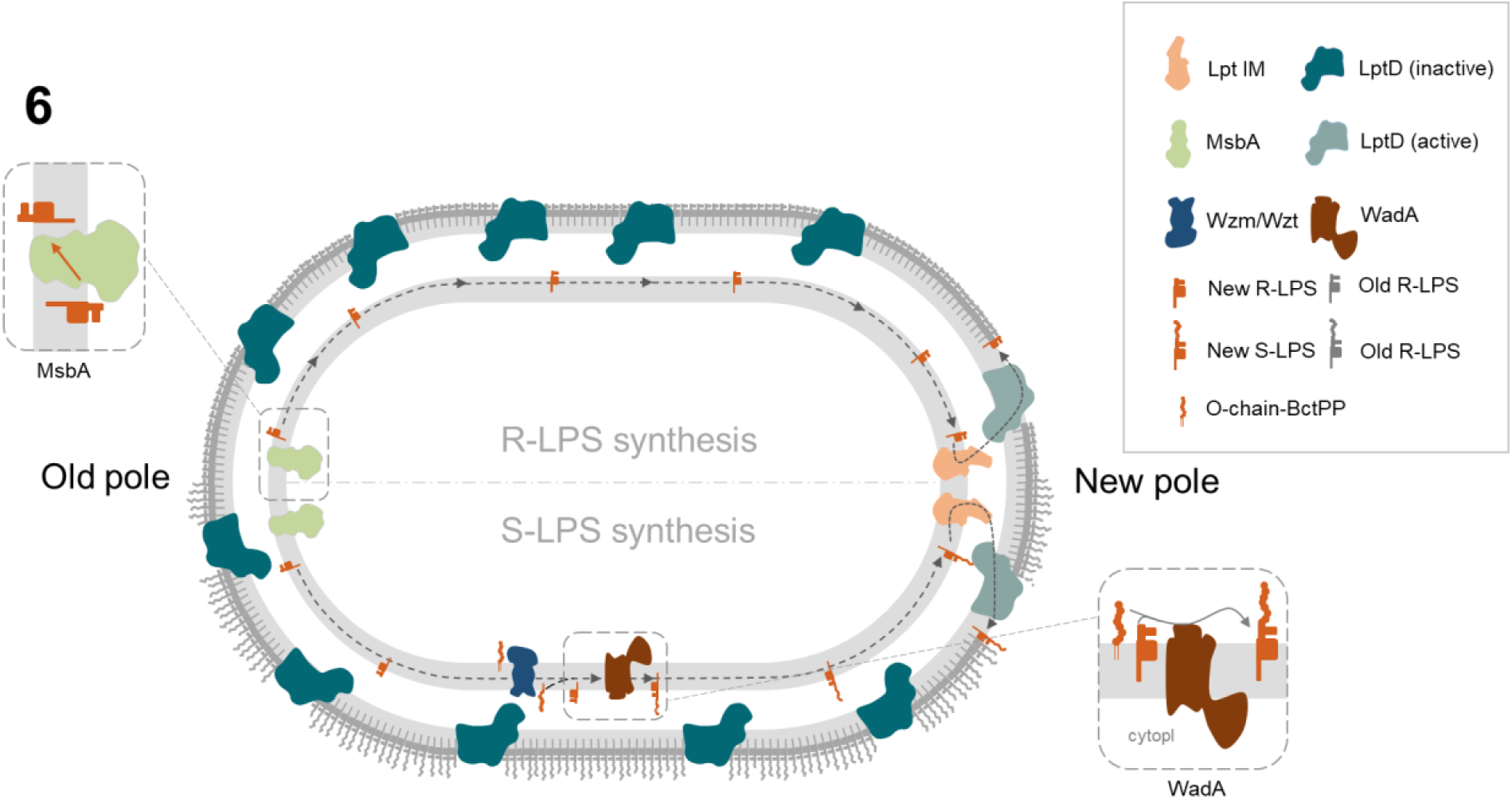
Schematic model localizing different actors involved in LPS synthesis and translocation pathway of *B. abortus*. When synthesized, the R-LPS is anchored to the inner leaflet of the IM. Then, the essential ABC transporter MsbA mainly localized at the old pole flips the R-LPS to the outer leaflet of the IM. The R-LPS is able to diffuse within the IM to the active growth sites (here the new pole) where the LPS translocation pathway, consisting of LptC and LptF is mainly localized. The R-LPS is translocated from the outer leaflet of the IM to the outer leaflet of the OM by the IM Lpt complex, LptA and finally the LptDE translocon. In this model, LptD localized at the growth site (light blue LptD) would actively incorporate new LPS molecules into the outer leaflet of the OM, whereas remaining homogenously dispersed LptD previously incorporated at the OM would remain inactive (dark blue LptD). On the lower part of the model, R-LPS is synthesized, flipped by MsbA and freely diffuses in the IM to WadA. WadA ligates the O-antigen onto the terminal core sugar to form S-LPS, which would freely diffuse in the IM until it reaches the Lpt system in order to be exported to the OM. Cytopl indicates the cytoplasm.

Since MsbA is mainly localized at the old pole and LptD is dispersed in the OM, we propose that the entire LPS transport system is not restrained at the growth areas in *B. abortus*. The analysis of a non-functional variant of MsbA suggests that MsbA function is coupled to its localization. One attractive hypothesis would be that old pole localization is driven by the availability of MsbA substrates, with inactive MsbA being excluded from these sites. The old pole localization of MsbA could be explained by the localization of the proteins involved in lipidA-core synthesis, but this needs to be further investigated. A main flippase activity at the old pole implies that lipid A-core needs to be mobile in the outer leaflet of the IM, from the old pole to the new pole of the cell, in non-dividing bacteria (Figure 6). This hypothesis is consistent with previous observations in *E. coli* suggesting that the whole IM is relatively fluid since it could be mixed in about 50 seconds (Kumar *et al*., 2010). These data reveal an unexpected aspect of LPS biosynthesis, involving trajectories in the IM and specific localization of enzymes, flippases and transporters that would be interesting to investigate in other bacteria.

Previous WadA characterization in *B. melitensis* convincingly showed that it is responsible for the addition of a glucose as the terminal sugar of the core, *i*.*e*. the anchor for the O-antigen (Salvador-Bescos *et al*., 2018). Here we show that the C-terminal domain of WadA, which is predicted to be an integral IM part of the protein with several periplasmic loops (Fig. S6), functions also as an O-antigen ligase in *B. melitensis* and *B. abortus*. To the best of our knowledge, it is the first time that a bifunctional O-antigen ligase is identified. It would add the terminal sugar on the core of LPS at the cytosolic side of the IM, and subsequently ligate the O-antigen once LPS precursor has been flipped over the IM by MsbA. Such a dual activity would ensure the grafting of an appropriate sugar (*i*.*e*. glucose) on a conserved part of the core (3-deoxy-D-manno-oct-2-ulosonic acid, KDO) to subsequently attach the O-antigen, and thus facilitate the functionality of the O-antigen ligase after an event of horizontal gene transfer. Otherwise, the success of horizontal gene transfer would be limited by the presence of an appropriate substrate in the core of LPS. Although few WadA homologs were identified in available databases, which is consistent with the horizontal gene transfer hypothesis, this bifunctional action mode could be found in other O-antigen ligases as well. WadA homologs display conserved residues, including some positively charged residues in the predicted periplasmic loop, and our data suggest that Arg614 plays an important role in the O-antigen ligase activity. WadA is thus forming a new class of O-antigen ligases, probably distantly related to previously described O-antigen ligases since WadA could be identified with a low similarity to existing O-antigen ligases.

In conclusion, this work highlights a new layer of spatiotemporal regulation of proteins involved in LPS synthesis and transport. Moreover, it reveals a new class of bifunctional O-antigen ligases. The investigation of homologous systems in other Gram-negative bacteria could be of great interest.

## Material and methods

### Bacterial strain and media

*Brucella abortus* 544 WT resistant to Nalidixic acid (Nal^R^) and derivative strains were grown in TSB rich medium (3% Bacto Tryptic Soy Broth) at 37°C. *Brucella melitensis* (Nal^R^) WT and derivative strains were grown in 2YT rich medium (1% yeast extract, 1.6% peptone, 0.5% NaCl) at 37°C. *Escherichia coli* DH10B and S17 strains were cultivated in LB medium at 37°C. All the strains used in this study are listed in Table S1.

When necessary antibiotics were added at the following concentrations; kanamycin (Kan, 10 or 50μg/ml, at the chromosomal locus or on a plasmid respectively), chloramphenicol (Cm, 4 or 20μg/ml), nalidixic acid (Nal, 25μg/ml) and Amp (100μg/ml).

### Strain construction

For deletions strains, about 500 base pairs upstream and downstream the region to be deleted were amplified and assembled by overlapping PCR. For Δ*OA-lig*, a major part of the main periplasmic loop was deleted, from amino acid 534 to 612. Translational fusions with the mNG were performed on the N-terminal or C-terminal part of the protein depending on the construct. Briefly, about 500 base pairs upstream and downstream the region of interest and the mNG gene were amplified by PCR. Except for RgsE and RgsS that were constructed using the Gibson assembly (see below), all the constructs were made by overlapping PCR, restriction and ligation in the corresponding plasmid. Apart from for RgsS and RgsE, a 3Flag tag was inserted downstream the mNG start codon. For LptC, the poorly conserved N-terminal part of the protein, TADIPHIHVP, was duplicated and used as a linker. For all the other constructs, RSATGS was used as a linker between the gene of interest and the mNG. Fusion and deletion strains were constructed by allelic replacement on the chromosomal locus as previously described using a pNPTS138 integrative plasmid (Kan^R^) (Deghelt *et al*., 2014).The additional copies of MsbA-mNG and MsbA_E491A_-mNG were provided on a pSK plasmid (Kan^R^), and was integrated on the genome via homologous recombination. Complementation strains were constructed using a pBBR1 MCS replicative plasmid carrying a chloramphenicol resistance cassette and the genes of interest were under the control of their endogenous promotor. In particular, for the complementation of the *OAg-lig* domain, aa from 267 to 703 were amplified and placed under the control of their endogenous promotor. For the induction of *gmd* expression a pBBR containing *gmd* under the control of the P_*lac*_ promoter was used, as previously described (Vassen *et al*., 2019). If required, the old pole marker or the new pole marker were integrated on a pSK (Kan^R^) or pKS(Cm^R^) respectively. All the primers and plasmids and ORFs used in this study are listed in Table S2, Table S3 and Table S4 respectively.

### Gibson assembly

Plasmid and inserts were amplified by PCR using primers with 20 overlapping base pairs designed with Benchling. The Gibson reaction was assembled on ice with 10μl of the 2x Gibson mix to which the plasmid (100ng) and inserts were added in a 1:1:1 ratio. Then, the reaction was kept at RT for 30 seconds and incubated overnight at 50°C. The 2X Gibson mix was made *in house*. For 800μl final volume, 0.6 μl T5 exonuclease (10 U/μl, NEB), 20 μl Phusion DNA polymerase (2 U/μl, NEB), 160 μl *Taq* DNA ligase (40 U/μl, NEB), 320 μl 5x reaction buffer (see below) and 300 μl of water. The 5x reaction buffer consisted of Tris-HCl 0.5 M, MgCl_2_ 50 mM, 1mM of each dNTPs, DTT 50 mM, ¼ W/V PEG 8000, NAD 5mM and dH_2_O.

### Induction of S-LPS

For the inducible strain of *B. abortus* 544 Δ*gmd* P_*lac*_-*gmd*, Δ*gmd*Δ*wadA* P_*lac*_-*gmd* and Δ*gmd*Δ*OAg-lig* P_*lac*_-*gmd*, early exponential phase culture (OD_600_ 0,2) were split in two parts; in TSB supplemented with 1mM of IPTG (induced) and in TSB without IPTG (non-induced). The cultures were incubated for 24h at 37°C under shaking. Then, bacteria were labelled with mAb against S-LPS (see below) and analyzed by fluorescence microscopy.

### IF

The LPS was visualized by IF using specific mAb against the O-antigen (A76/12G12). Exponential phase cultures were washed 2 times in PBS at 7000 rotations per minute (r.p.m). for 2.5 min, and resuspended in the supernatant for the hybridoma culture containing the mAb. After 40 min of incubation at RT on a rotating wheel, the samples were washed 3 times in PBS and the pellet was resuspended in the secondary anti mouse antibody coupled to AlexaFluor 544 (1:500, in PBS), and incubated at RT on the rotating wheel for 1h. Bacteria were then washed 3 times, resuspended in PBS and analyzed by fluorescence microscopy.

### eFluor labelling

Bacteria in early exponential phase (OD_600_ 0.2) were washed twice in phosphate buffered saline (PBS) and resuspended in eBioscience™ Cell Proliferation Dye eFluor™ 670 (eFluor, Invitrogen) at a final concentration of 1 μg/ml in PBS. After 15 min of incubation protected from light, bacteria were washed 3 times, resuspended in preheated TSB medium. After 3h of incubation at 37°C under constant agitation, the bacteria were washed 2 times and resuspend in PBS and analysed by fluorescence microscopy.

### Microscopy and analysis

Exponential phase bacterial cultures were washed two times and resuspended in PBS. Two μl of the bacterial suspension were spotted onto 1% agarose pads. Images were acquired using a Nikon Eclipse Ti2 equipped with a phase-contrast objective Plan Apo λ DM100XK 1.45/0.13 PH3 and a Hamamatsu C13440-20CU ORCA-FLASH 4.0. Fluorescence images were analysed using FIJI v.2.0.0 (Schindelin *et al*., 2012), a distribution of ImageJ. Look-up tables (LUT) were adjusted to the best signal–background ratio. For demograph analysis, bacteria were detected and analysed using MicrobeJ, a pluggin of ImageJ (Ducret *et al*., 2016), a plugin of the ImageJ software. Only isolated bacteria were taken into account, aggregates were excluded from the analysis. If necessary, small aggregates or incorrectly selected bacteria were manually removed from the analysis. Whether the strains contained the old (PdhS-mCherry) or new pole marker (PopZ-mCherry), only the bacteria containing one focus were retained for the analysis. Due to technical variability (*e*.*g*. thickness of the agarose pad), different time of exposure, LUT, tolerance for focus detection and minimal fluorescence intensity were used for the different analysis.

### Western blot analysis

Exponential phase cultures were normalised to OD_600_ 10 in PBS and inactivated for 1h at 80°C. SDS-β-mercaptoethanol loading buffer was added at 1:4 of final volume. Samples were heated at 95 °C for 15 min and were loaded on 12% acrylamide gels. After migration, proteins and LPS were transferred onto a nitrocellulose membrane (GE Healthcare Amersham Protran 0.45 NC) and blocked overnight at 4 °C in PBS supplemented with 0.05% Tween-20 (VWR) and 5% (w/v) milk (Nestlé, Foam topping). Membranes were washed three times for 10 min in PBS 0.05% Tween-20. Afterwards, membranes were incubated for 1 h at room temperature (RT) with primary antibodies (anti-core-LPS A68/24D08/G09 and anti-3Flag FG4R (Thermo Fisher Scientific), 1:100; anti-LptD, 1:100) (Table S4). Membranes were then incubated for 1h at RT with the corresponding secondary horseradish-peroxidase-coupled antibody (1:5000, Table S5). Both antibodies were diluted in PBS supplemented with 0.05% Tween-20 and 0.5% milk, and the membranes were washed as described after both incubations. The membranes were revealed using Clarity ECL Substrate (Bio-Rad) solutions and images were acquired using GE Healthcare Amersham Imager 600.

### LptD antibody production

In order to generate polyclonal Ab against LptD for Western blot analysis, the predicted N-terminal soluble part of LptD without signal peptide (nucleotides 109-666 bp from *B. abortus lptD*) was amplified by PCR and cloned into the overexpression vector pET15b generating a N-terminal His6 tag fusion (pET15b_lptD). Overexpression was done in *E. coli* BL21(DE3) in the presence of carbenicillin. Bacterial culture was induced with IPTG (1 mM final concentration) for 4 h at 37 °C shaking, lysed by sonication and purified with Ni-NTA resin. The insoluble fraction was further treated in the presence of urea. The soluble and insoluble fractions were analyzed by SDS-PAGE. The protein was present in the insoluble fraction and used for immunization in a rabbit. 50 μg of antigens with Complete Freund Adjuvant were injected at day 0. Every 4 weeks, 50 μg of antigens with Incomplete Freund Adjuvant were injected until 5 injections in total. Samples of bleeding were taken after 84, 112 and after 120 days for final bleeding. The antisera were tested by Western blot and used at a final dilution of 1:5000.

### IF- Scanning electron microscopy (IF-SEM)

Eighty μl of exponential growth phase bacteria were washed twice with PBS (7000 r.p.m., 2.5 min) and fixed with 2% paraformaldehyde (PFA) for 20 min at RT. After two washings with PBS, bacteria were labeled with an Ab directed against the 3Flag (1:20) (Thermo Fisher Scientific) and a secondary Ab anti-mouse IgG conjugated with 18 nm gold particles (1:10) (Abcam). Fifty μg/ml of poly-L-lysine (Sigma-Aldrich) were incubated for 1h at RT on a coverslip, bacteria were added on coated coverslips and centrifuged at 2000 r.p.m. for 5 min. Samples were post-fixed with 200 μl of 2.5% glutaraldehyde in 0.1 M cacodylate buffer pH 7.4 for 1h at RT. After two washings with 0.2 M cacodylate buffer pH 7.4, cover slips were dehydrated in ethanol with increasing concentrations from 30% to 100% ethanol. Critical Point Drying was done with CPD030 critical point dryer (BALZER) and samples were stored air-sealed at RT upon observation. Immediately before imaging, samples were coated with 5 nm chromium (Quorum Q150T ES, Quorum Technologies) and observed with the field emission scanning electron microscope JEOL JSM-7500 F. Images were acquired with the following settings: 20 μA emission current, 15 kV accelerating voltage and probe current of 9. Isolated bacteria were imaged with a magnification of 40,000x by secondary electron imaging (SEI) and subsequently by detection of low angle backscatter electrons (LABE mode) to visualize the gold particles on the bacterial surface. Further image analysis was carried out with ImageJ (Schneider *et al*., 2012).

### SEM image analysis

Images recorded by SEM were analyzed with ImageJ (Schneider *et al*., 2012) as followed. For non-dividing bacteria, the length was measured with a straight line and bacteria were divided in the middle in two equal parts by a line at a right angle to the main axis of the bacterium (Fig. 51a). From this midcell line, a straight line measured the distance from the midcell to the gold particle. Bacteria showing a visible constriction site were classified as dividing bacteria. If dividing bacteria were found in nearly straight position, bacteria were divided at the constriction site and the distance of gold articles to the constriction site was measured (Fig. 51b). If dividing bacteria displayed a curved position, the lengths of both cells (mother and future daughter cell) were measured first by straight lines between both pole and constriction site (Fig. 51c). From this cell length line, a straight line in 90° angle was set to the pole facing the constriction site. Distances were measured from the center of the gold particles towards the line highlighting the constriction site. The ratio between distance of gold particles and length of half bacterium or constriction site, respectively, was calculated for each detected gold particle and results were summarized in 5 categories ranging from 0 (close to the middle or constriction site) to 1 (close to the poles). Given that these measurements were performed on three independent samples, distribution of gold particles on the bacterial surface was statistically analyzed by t-test in comparison to the theoretical frequency of random distribution (20%). In order to analyze the asymmetry between the two sides of the bacterium, the standard deviation for a random insertion was calculated using a binomial distribution based on a probability of occurrence of 0.5 for each side of the bacterium. If the difference of particle number between the two sides of the bacterium exceeds 2 standard deviations, it was considered that particles distribution was asymmetric. To analyze the asymmetry between cell poles of non-dividing bacteria, bacteria ≤ 1 μm were analyzed.

## Acknowledgements

We are grateful to Anke Becker and Elizaveta Kroll for the discussion and help regarding RgsS and RgsE. We thank the Electron microscopy facility of UNamur. We thank Raquel Conde-Alvarez and Jean Michel Jault, Jean-François Collet and Jean-Yves Matroule for for their insightful discussions and advices, as well as Adrien Ducret for his help with MicrobeJ. We also thank Neeraj Dhar for advices regarding mNeonGreen and Mathieu Waroquier and Françoise Tilquin for their technical assistance. This research has been funded by grants from the *Fonds de la Recherche Scientifique-Fonds National de la Recherche Scientifique* (FRS-FNRS, http://www.fnrs.be) (PDR Brucell-cycle T.0060.15, and PDR Single cell analysis of *Brucella* growth T.0058.20) to X. De Bolle. The work was also supported by a grant from Actions de Recherche Concertée 17/22-087 from the *Fédération Wallonie-Bruxelles* of Belgium. V. Vassen was founded by a PhD fellowship from FRIA (FNRS).

