## Supplemental tables and figures for "Lipopolysaccharide synthesis and traffic in the envelope of the pathogen *Brucella abortus*"

**Table S1| List of strains used in this study.**

**Table S2 | List of primers used in this study.**

**Table S3 | List of plasmids used in this study**

**Table S4 | List of primary antibodies used in this study.**

**Table S5 | List of secondary antibodies used in this study.**

**Table S6 | Table of ORF investigated in this study.**

**Figure S1 | Alignment of Rhizobiales LptD amino acid sequences.**

**Figure S2 | Localization of LptD.**

**Figure S3 | Growth behaviour of *mNG-lptC*, *mNG-lptF* and *msbA-mNG* strains.**

**Figure S4 | MsbA localization in *B. abortus.***

**Figure S5 | S-LPS phenotype of *B. abortus* Δ*waaL.***

**Figure S6 | Sequence alignment and structure prediction for WadA as the O-antigen ligase in *B. abortus.***

**Figure S7 | S-LPS labelling of *B. melitensis* 16M strains.**

**Table S3| List of strains used in this study.**

| Name | Genotype | Reference |
|---|---|---|
| | *Brucella abortus* strains | |
| **Wild type (WT)** | *B. abortus* 544, $Nal^R$ | J.-M.Verger, INRA, Tours |
| ***rgsS-mNG pdhS-mCherry*** | *B. abortus* 544 *rgsS-mNG pdhS*::pSK kan *pdhs-mCherry* | This study |
| ***rgsE-mNG pdhS-mCherry*** | *B. abortus* 544 *rgsE-mNG pdhS*::pSK kan *pdhs-mCherry* | This study |
| ***lptD-AGDP*** | *B. abortus* 544 *lptD-AGDP-G4-3Flag* | V. Vassen, PhD thesis, UNamur, BE |
| ***mNG-lptC pdhS-mCherry*** | *B. abortus* 544 *mNG-lptC pdhS*::pSK kan *pdhs-mCherry* | This study |
| ***mNG-lptF pdhS-mCherry*** | *B. abortus* 544 *mNG-lptF pdhS*::pSK kan *pdhs-mCherry* | This study |
| ***msbA-mNGpdhS-mCherry*** | *B. abortus* 544 *msbA-mNG pdhS*::pSK kan *pdhs-mCherry* | This study |
| **pSK_*msbAmNG popZ-mCherry*** | *B. abortus 544 msbA::pSK kan msbA-mNG popZ::pSK kan popZ-mCherry* | This study |
| **pSK_*$msbA_{E491A}$-mNG popZ-mCherry*** | *B. abortus* 544 *msbA*::pSK kan *$msbA_{E491A}$-mNG popZ*::pSK kan *popZ-mCherry* | This study |
| ***Δgmd*** | *B. abortus* 544 Δ*gmd* | (Vassen *et al.*, 2019) |
| ***ΔwaaL*** | *B. abortus* 544 Δ*waaL* | This study |
| ***ΔgmdΔwadA*** | *B. abortus* 544 Δ*gmd*Δ*wadA* | This study |
| ***ΔgmdΔOAg-lig*** | *B. abortus* 544 Δ*gmd*Δ*OAg-lig* | This study |
| ***Δgmd* pBBRI *gmd*** | *B. abortus* 544 Δ*gmd* pBBRI_*gmd* | This study |
| ***ΔgmdΔwadA* pBBRI *gmd*** | *B. abortus* 544Δ*gmd*Δ*wadA* pBBRI_*gmd* | This study |
| ***ΔgmdΔOAg-lig* pBBRI *gmd*** | *B. abortus* 544 Δ*gmd*Δ*OAg-lig* pBBRI_*gmd* | This study |
| | *Brucella melitensis* 16M | |
| **Wild type (WT)** | *B. melitensis* 16M, $Nal^R$ | A. Macmillan, Central Veterinary Laboratory, Weybridge, UK |
| ***Δgmd*** | *B. melitensis* 16M Δ*gmd* | This study |
| ***ΔwadA*** | *B. melitensis* 16M Δ*wadA* | This study |
| ***ΔOAg-lig*** | *B. melitensis* 16M Δ*OAg-lig* | This study |
| ***ΔwadA* pBBR *wadA*** | *B. melitensis* 16M Δ*wadA* pBBR_MCSI_*wadA* | This study |
| ***ΔOAg-lig* pBBR *OAg-lig*** | *B. melitensis* 16M Δ*OAg-lig* pBBR_MCSI_*OAg-lig* | This study |
| ***ΔwadA* pBBR *wadA-R614A*** | *B. melitensis* 16M Δ*wadA* pBBR_MCSI_*wadA-R614A* | This study |

**Table S4 | List of primers used in this study.** Nucleotides not annealing on DNA template are shown in bold, red characters represent point mutations and UPPERCASE corresponds to restriction site.

| Primer name | 5'-3' sequence |
|---|---|
| **pNPTS138-rgsSmNG-F** | **ttccaaaaacccccatgcgg**atctggatccacgaattcgc |
| **pNPTS138-rgsSmNG-R** | **ttgcgcattgttctgcggta**atcctgcagagaagcttggc |
| **rgsSmNG-UP-F** | **gccaagcttctctgcaggat**taccgcagaacaatgcgcaa |
| **rgsSmNG-UP-R** | **acggatcctgttgcagaacg**ctgcgtcacgaaacagcttc |
| **mNG-4-rgsS- F** | **agcgttctgcaacaggatcc**gtctcgaagggcgaagaaga |
| **mNG-4-rgsS- R** | **tggaaccttgctggttgttt**tcacttatacagttcatccatgc |
| **rgsSmNG-DW-F** | **tggatgaactgtataagtga**aaacaaccagcaaggttcca |
| **rgsSmNG-DW-R** | **gcgaattcgtggatccagat**ccgcatgggggtttttggaa |
| **check-rgsSmNG-F** | caagtgttgccggtgtg |
| **check-rgsSmNG-R** | tcattgagcgccttgaacg |
| **pNPTS138-rgsEmNG-F** | **gagcccggcgcagagctgac**atctggatccacgaattcgc |
| **pNPTS138-rgsEmNG-R** | **ttcagcaagttccttgagcg**atcctgcagagaagcttggc |
| **rgsEmNG-UP-F** | **gccaagcttctctgcaggat**cgctcaaggaacttgctgaaattgt |
| **rgsEmNG-UP-R** | **tcttcttcgcccttcgagac**ccgaaggcgtccgcttgcat |
| **mNG-4-rgsE-F** | **atgcaagcggacgccttcgg**gtctcgaagggcgaagaaga |
| **mNG-4-rgsE-R** | **ttttaactggagccgctcta**cttatacagttcatccatgccc |
| **rgsEmNG-DW-F** | **gcatggatgaactgtataag**tagagcggctccagttaaaattgagca |
| **rgsEmNG-DW-R** | **gcgaattcgtggatccagat**gtcagctctgcgccgggctc |
| **check-rgsEmNG-F** | aagcgaaggaaacgacgac |
| **check-rgsEmNG-R** | acttgcaattgctgatccc |
| **lptD_AGDP_UP_F_SpeI** | **ctaga**ACTAGTgaaatcaccacattcaacaatgg |
| **lptD_AGDP-G4-3F-4_UP-R2** | **cgatgtcatgatctttataatcaccgtcatggtctttgtagtcgccaccgccgcc**cgg atcgcccgccaggtag |
| ***lptD*_AGDP-G4-3F-G4_DW-F** | **ttataaagatcatgacatcgattacaaggatgacgatgacaagggcggcggtggc**aatga aatgtattcaaagcagc |
| ***lptD*_AGDP_DW-R_SalI** | **tgcac**GTCGACgccaagctggaaggactg |
| **mNGlptC-UP-F** | caggatgacacccttcac |
| **mNGlptC-UP-R** | **tcatggtctttgtagtccat**gttatcctttggcacatca |
| **mNG-4-lptC-F** | atggactacaaagaccatga |
| **mNG-4-lptC-R** | cggaacatgaatatgtggta |
| **mNGlptC-DW-F** | **taccacatattcatgttccg**acggccgacataccac |
| **mNGlptC-DW-R** | ttgtttgccgaaagcgtg |
| **check-mNGlptC-F** | aaaagggtcatgccacg |
| **check-mNGlptC-R** | tcgtgcatgtctttatccc |
| **mNGlptF-UP-F** | ggcagtcttgcccaaag |
| **mNGlptF-UP-R** | **gtcatggtctttgtagtccat**aggtggagtctacatccc |
| **mNG-4-lptF-F** | atggactacaaagaccatga |
| **mNG-4-lptF-R** | **aaaTCTAGA**cttatacagttcatccatgc |
| **mNGlptF-DW-F** | **aaaTCTAGAcggtcggccaccggatcc**cgattgatcgagttgtaca |

| | |
|---|---|
| **mNGlptF-DW-R** | ttttccatcagcagcatg |
| **check-mNGlptF-F** | ttcctctctcatgtgaga |
| **check-mNGlptF-R** | ccgaatcctgattcatcg |
| **msbAmNG-UP-F** | ttctatgacctcgaccgg |
| **msbAmNG-UP-R** | **ggatccggtggccgaccg**tctggcctctcctgtg |
| **mNG-4-msbAm-F2** | **cggtcggccaccggatcc**gactacaaagaccatgacg |
| **mNG-4-msbAm-R2** | **cacttcctcgctca**cttatacagttcatccatgcc |
| **msbAmNG-DW-F** | tgagcgaggaagtgcaa |
| **msbAmNG-DW-R** | ggccagtatagccatcac |
| **check-msbAmNG-F** | tgcgcaacctctatctg |
| **check-msbAmNG-R** | ggccagtatagccatcac |
| **pSKmsbAmNG-F** | **aaaACTAGT**tgcgcaacctctatctg |
| **pSKmsbAmNG-R** | **ttttTCTAGA**tcacttatacagttcatccatg |
| **pSKmsbA**$_{E491A}$**-mNG-UP-F** | **aaaACTAGT**tgcgcaacctctatctggg |
| **pSKmsbA**$_{E491A}$**-mNG-UP-R** | **ttgtcgagcgcggacgttgctg**cgtcgagaagcaggaccg |
| **pSKmsbA**$_{E491A}$**-mNG-DW-F** | tcgacg**ca**gc**a**acgtcc |
| **pSK msbA**$_{E491A}$**-mNG-DW-R** | **tttTCTAGA**tcacttatacagttcatccatgccc |
| **check-Δgmd-F** | agctaacttgctggcataag |
| **check-Δgmd R** | caggtcgagtcgatcatg |
| **ΔwaaL-UP-F** | aattgcaatgcctataatgc |
| **ΔwaaL-UP-R** | **ttcatgaagtcactccga**tcagatatccaaaatgtgcc |
| **ΔwaaL-DW-F** | tcggagtgacttcatgaa |
| **ΔwaaL-DW-R** | cgccatatgcagatgttcg |
| **ΔwadA-UP-F** | gatcaagcaggtgctcatc |
| **ΔwadA-UP-R** | **tgtaaaagtggaaccgctc**taattcgcttgcctcagc |
| **ΔwadA-DW-F** | gagcggttccacttttaca |
| **ΔwadA-DW-R** | gcaaatatcagcgcaacg |
| **check-ΔwadA-F** | gaaacggtgcggataatc |
| **check-ΔwadA-R** | aacgcatatcgaagcttt |
| **ΔOAg-lig-UP-F** | cgaggttcgcatatatctatc |
| **ΔOAg-lig-UP-R** | taacaattcccaggatgtatt |
| **ΔOAg-lig-DW-F** | **aatacatcctgggaattgtta**attcgctatggatttctgg |
| **ΔOAg-lig-DW-R** | gaagtgctggcaaacttt |
| **check-ΔOAg-lig-F** | gaaacggtgcggataatc |
| **check-ΔOAg-lig-R** | aacgcatatcgaagcttt |
| **pBBRI-wadAc-F** | **tttCTGCAG**atttctcgctcgatcgc |
| **pBBRI-wadAc-R** | **tttCTCGAG**attacaggcttgtcaggg |
| **pBBRI-OAg-lig-UP-F** | **ttttGAGCTC**atttctcgctcgatcgc |
| **pBBRI-OAg-lig-UP-R** | **aaaaCATATG**cattaattcgcttgcctca |
| **pBBRI-OAg-lig-DW-F** | **aaaCATATG**tccgatcagaattcaggtt |
| **pBBRI-OAg-lig-DW-R** | **aaaaCTGCAG**attacaggcttgtcaggg |

| | |
|---|---|
| **pBBRwadA-R614A-UP-F** | **tttCTGCAG**atttctcgctcgatcgc |
| **pBBRwadA-R614A-UP-R** | catag**gc**aatggcaatttccag |
| **pBBRwadA-R614A-DW-F** | **ctggaaattgccattgcctatg**gatttctgggcttgctgt |
| **pBBRwadA-R614A-DW-F** | **tttCTCGAG**attacaggcttgtcaggg |

**Table S3 | List of plasmids used in this study**

| Plasmid name | Reference |
|---|---|
| **pNPTS138** | M. R. K. Alley, Imperial College of Science,London, UK |
| **pSKoriT kan** | (Haine *et al.*, 2005) |
| **pKSoriT cat** | Isabelle Danese and Pascal Lestrate, UNamur, BE |
| **pBBR_MCSI** | (Kovach *et al.*, 1994) |
| **pNPTS138_*rgsS-mNG*** | This study |
| **pNPTS138_*rgsE-mNG*** | This study |
| **pNPTS138 *lptD::lptD-3Flag*** | V. Vassen, PhD thesis, UNamur, BE |
| **pNPTS138_*mNG-lptC*** | This study |
| **pNPTS138_*mNG-lptF*** | This study |
| **pNPTS138_*msbA-mNG*** | This study |
| **pSKoriT_kan_*pdhS-mCherry*** | (Van der Henst *et al.*, 2010) |
| **pKSoriT_cat_*popZ-mCherry*** | (Deghelt *et al.*, 2014) |
| **pSKoriT_kan-*msbA-mNG*** | This study |
| **pSKoriT_kan- *msbA*$_{E491A}$*-mNG*** | This study |
| **pNPTS138_Δ*waaL*** | This study |
| **pNPTS138_Δ*wadA*** | This study |
| **pNPTS138_Δ*OAg-lig*** | This study |
| **pBBR_MCSI_*waaL*** | This study |
| **pBBR_MCSI_*wadA*** | This study |
| **pBBR_MCSI_*OAg-lig*** | This study |
| **pBBR_MCSI_*wadA-R614A*** | This study |
| **pBBRI_P$_{lacZ}$_*gmd*** | (Vassen *et al.*, 2019) |
| **pNPTS138_Δ*gmd*** | (Vassen *et al.*, 2019) |

**Table S4 | List of primary antibodies used in this study.**

| Name | Recognized structure | Host (Isotype) | Application | Reference |
|---|---|---|---|---|
| **A76/12G12** | *Brucella* S-LPS (O-antigen) | Mouse (IgG1) | WB, IF | (Cloeckaert *et al.*, 1993) |
| **A68/24D08/G09** | *Brucella* R-LPS (lateral branch of the core) | Mouse (Unknown) | WB | (Bowden *et al.*, 1995) |
| **Anti-LptD** | *Brucella* LptD | Rabbit (Polyclonal) | WB | This study |
| **DYKDDDDK Tag (FG4R)** | 3Flag tag | Mouse (IgG2b) | WB, IF- SEM | ThermoFisher Scientific |
| **A68/7G11/C10** | *Brucella* Omp10 | Mouse (IgG2a) | WB | (Cloeckaert *et al.*, 1990) |

**Table S5 | List of secondary antibodies used in this study.**

| Target | Conjugate | Application | Reference |
|---|---|---|---|
| **Mouse IgG (H+L)** | Alexa Fluor 514 | IF | Invitrogen (A-31555) |
| **Mouse** | HRP | WB | Dako |
| **Rabbit** | HRP | WB | Dako |
| **Mouse** | 18 nm gold | IF-SEM | Abcam |

**Table S6 | Table of ORF investigated in this study**

| Gene name | ORF number in *B. abortus* 2308 | Uniprot accession number |
|---|---|---|
| *rgsS* | BAB1_0897 | Q2YNJ8 |
| *rgsE* | BAB1_1043 | Q2YQ47 |
| *lptD* | BAB1_0707 | Q2YN48 |
| *lptC* | BAB1_0154 | Q2YP16 |
| *lptF* | BAB1_0709 | Q2YN46 |
| *msbA* | BAB2_1011 | Q2YJN8 |
| *gmd* | BAB1_0545 | Q2YMP3 |
| *waaL* | BAB2_0106 | Q2YJ92 |
| *wadA* | BAB1_0639 | Q2YMX2 |

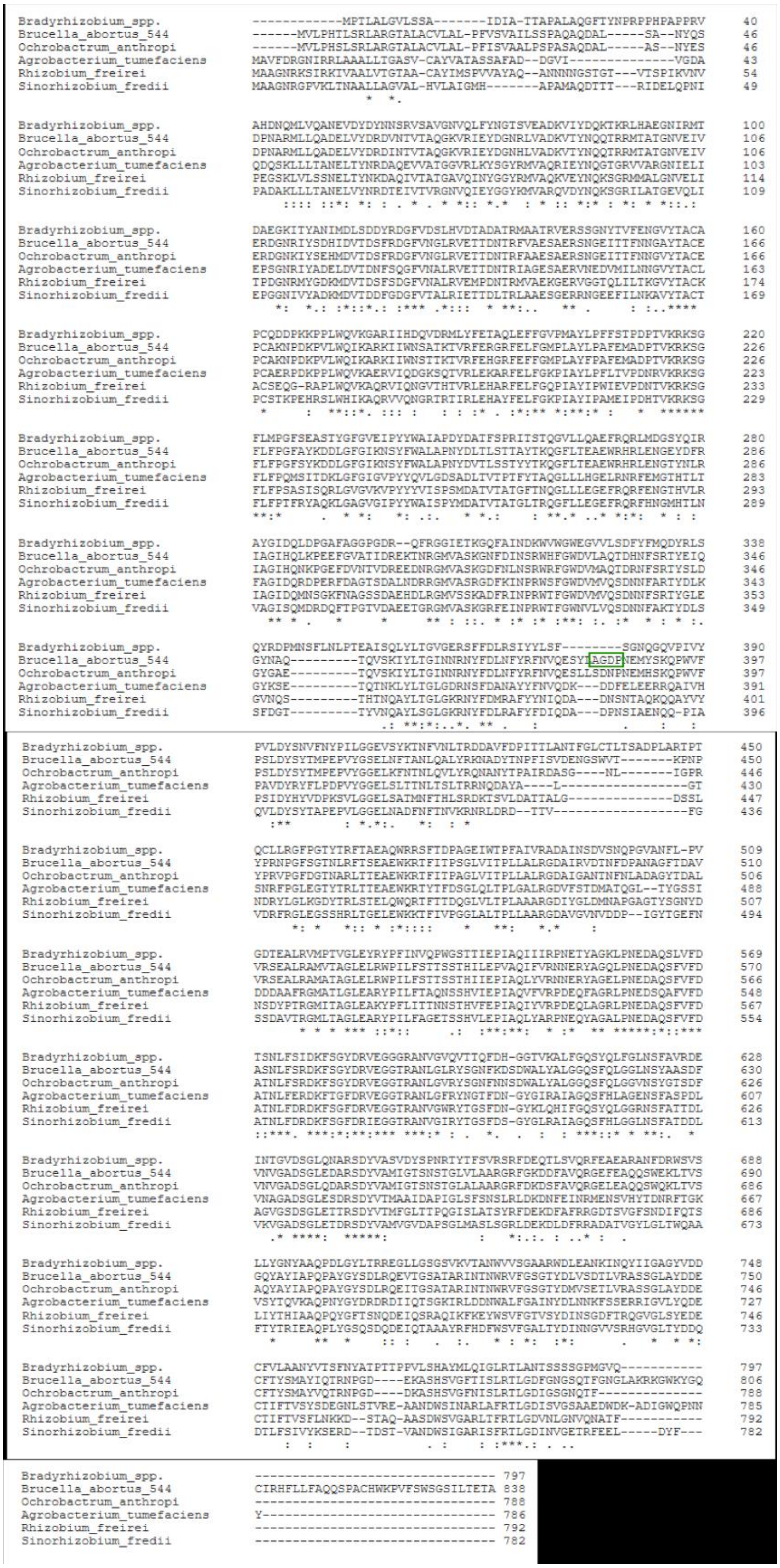

**Figure S1 | Alignment of Rhizobiales LptD amino acid sequences.** Sequences of LptD homologs in other Rhizobiales were identified by blastp (Altschul *et al.*, 1997) and multiple sequence alignment was generated with CLUSTAL Omega (1.2.4) (Sievers *et al.*, 2011). Position of the insertion of a 3Flag tag is shown with a green box (AGDP loop, position 383 in *B. abortus* 544 sequence). The ADGP sequence was duplicated and a $Gly_4$ linker sequence was inserted on each side of the 3Flag.

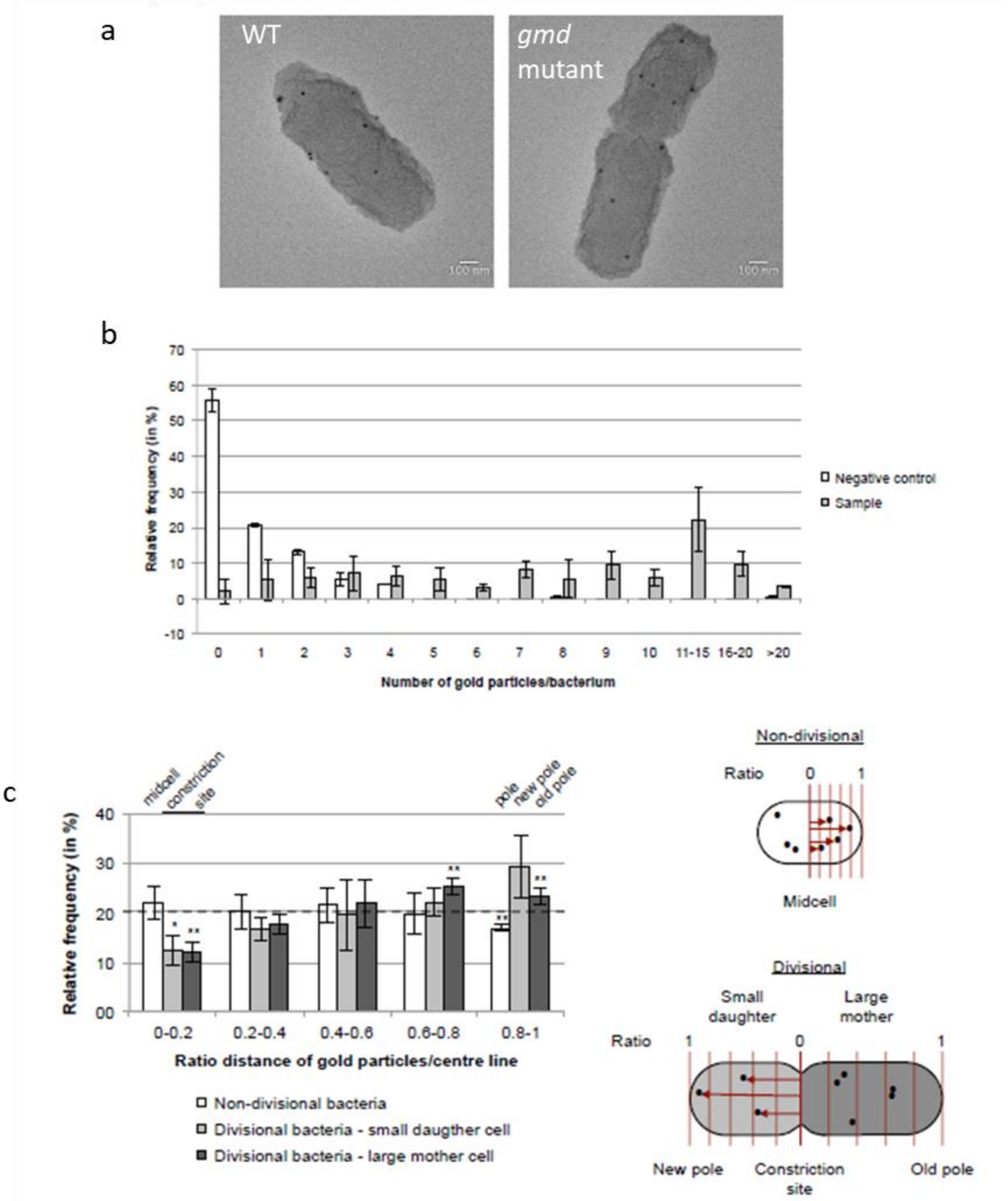


.

**Figure S2 | Localization of LptD.** a, SEM images from exponential phase culture of *B. abortus* WT and *3Flag::lptD*. Scale bar is 100 nm. Images were acquired by low angle backscattered electron (LABE) mode and inverted. Black dots correspond to gold particles. Brightness (+20%) and sharpness (+50%) of images were adjusted (b) Frequency distribution of gold particles from negative control (*B. abortus* disrupt *gmd*) and sample (*B. abortus* 544 disrupt *gmd lptD*-3Flag) after IF-SEM. Error bars corresponds to standard deviation from two and three independent experiments, respectively. $n_{negative\ control}$= 142 bacteria, $n_{sample}$=169 bacteria. **(c)** The distribution of gold particles labelling 3Flag::LptD was analyzed by measuring the distance towards central line in non-divisional and towards constriction site in divisional bacteria. Ratios of distance of gold particles/central line were classified in five categories (model on right side, ratio distances as shown as red arrows). Divisional bacteria were further divided in daughter and mother part according to their size. Black dashed line represents theoretical frequency of random distribution (20%). Statistical differences compared to relative frequency of 20% were analyzed by t-test. *$p$<0.05, **$p$<0.01. Error bars correspond to standard deviation from three independent experiments. Number of non-divisional bacteria= 58, Number of divisional bacteria=77.

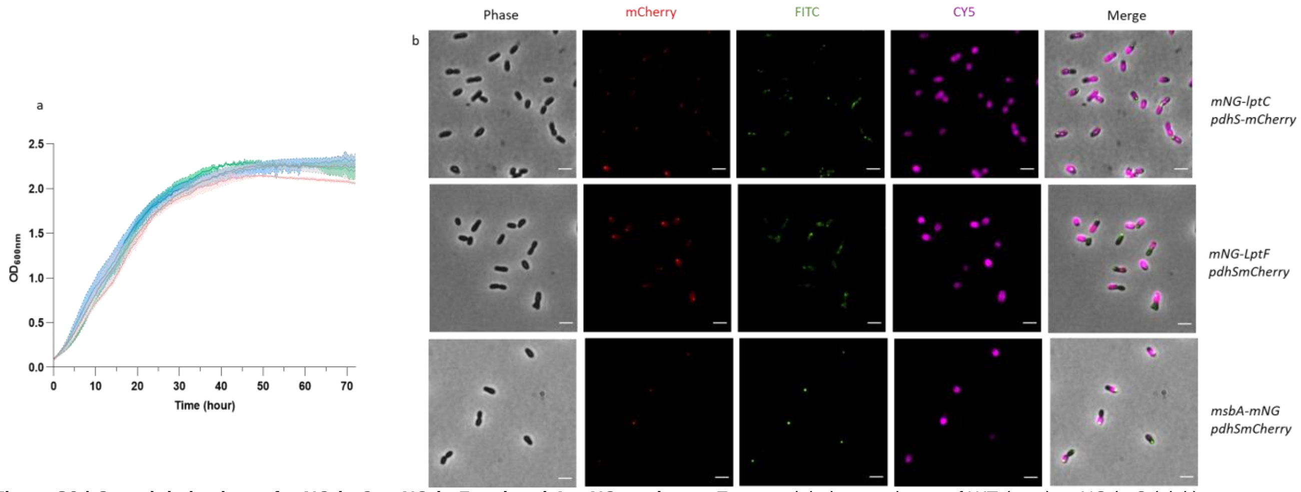


**Figure S3 | Growth behaviour of *mNG-lptC*, *mNG-lptF* and *msbA-mNG* strains. a,** Exponential phase cultures of WT (grey), *mNG-lptC* (pink), *mNG-lptF* (blue) and *msbA-mNG* (green) were diluted to $OD_{600}$ =0.1 and grown in TSB rich medium at 37°C for 72 h. The $OD_{600}$ was measured every 30 minutes. Dotted lines correspond to one standard deviation. N=3. **b,** eFluor labelling of *mNG-lptC*, *mNG-lptF* and *msbA-mNG* (all green) was performed on early exponential phase culture in strains co-expressing PdhS-mCherry. Bacteria were labelled with eFluor, washed and grown for 2.5 h at 37°C. The unlabelled part corresponds to newly incorporated material. Scale bars are 2 µm.

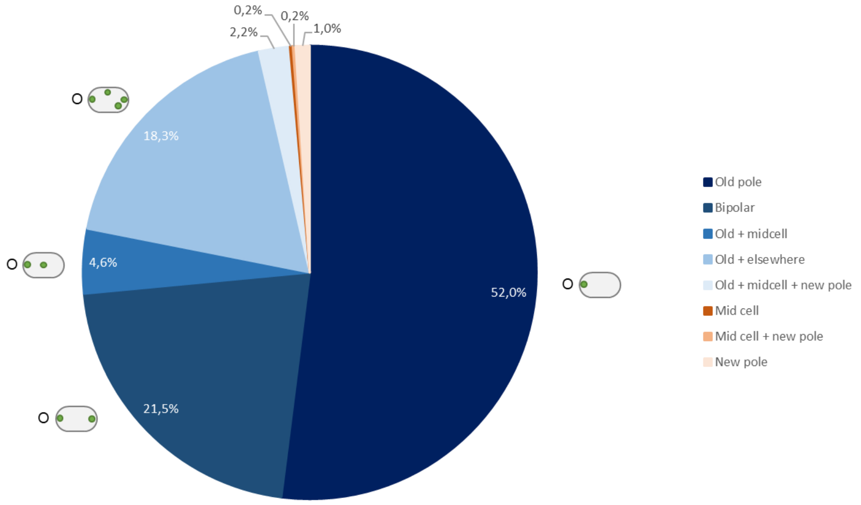


**Figure S4 | MsbA localization in *B. abortus*.** MsbA is found at the old pole in 98,6% of bacteria. 52% of the cells display only one focus of MsbA at the old pole, 21,5% show a bipolar localization of MsbA,4,6% display one focus at the mid cell and the old pole, and 18,3% show one focus of MsbA and at least another somewhere else in the bacterium. 2,2% percent of the bacteria display one focus at the old pole, mid cell and at the new pole. Only 0.2% had a focus at the mid cell, 0.2% at the new pole and the constriction site and 1% had one focus at the new pole only. The analysis was performed by manual counting of bacteria of the strain co-expressing MsbA-mNG and PdhS-mCherry labelled with eFluor. O: Old pole. N=540 bacteria.

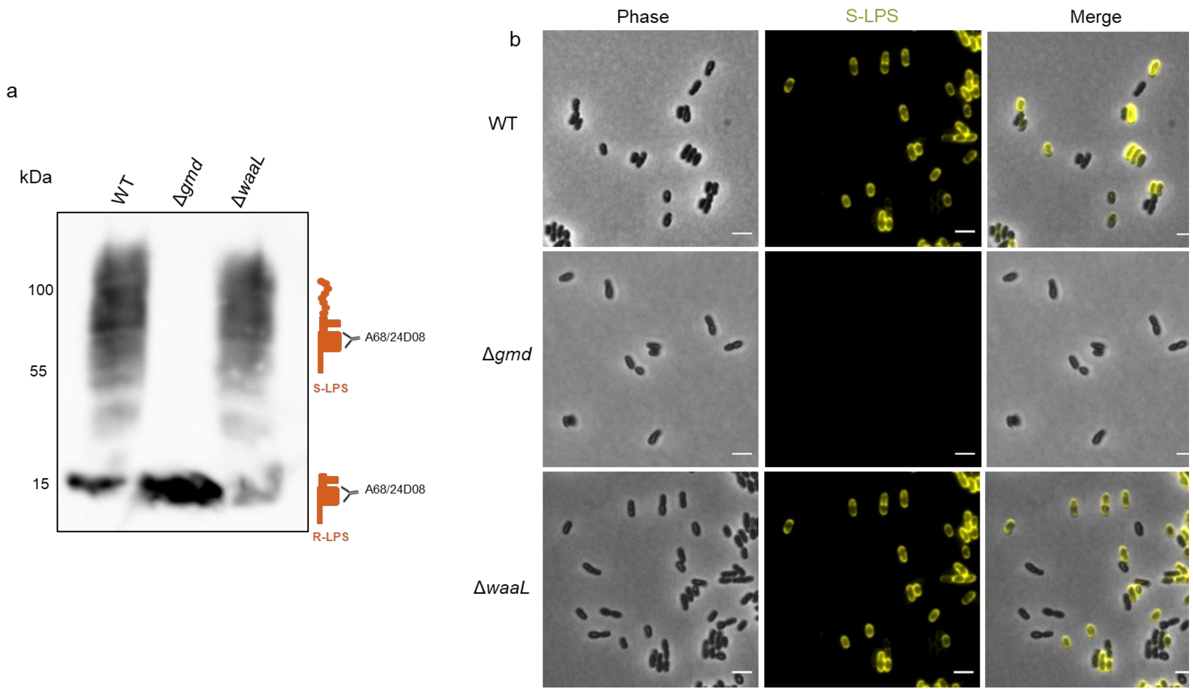


**Figure S5 | S-LPS phenotype of *B. abortus* Δ*waaL*.** a, WB analysis performed using the A68/24D08 mAb targeting the core of both R-LPS and S-LPS. b, Immunofluorescence microscopy labelling S-LPS of WT, Δ*gmd* and Δ*waaL* using the mAb A76/12G12. Scale bars are 2 µm.

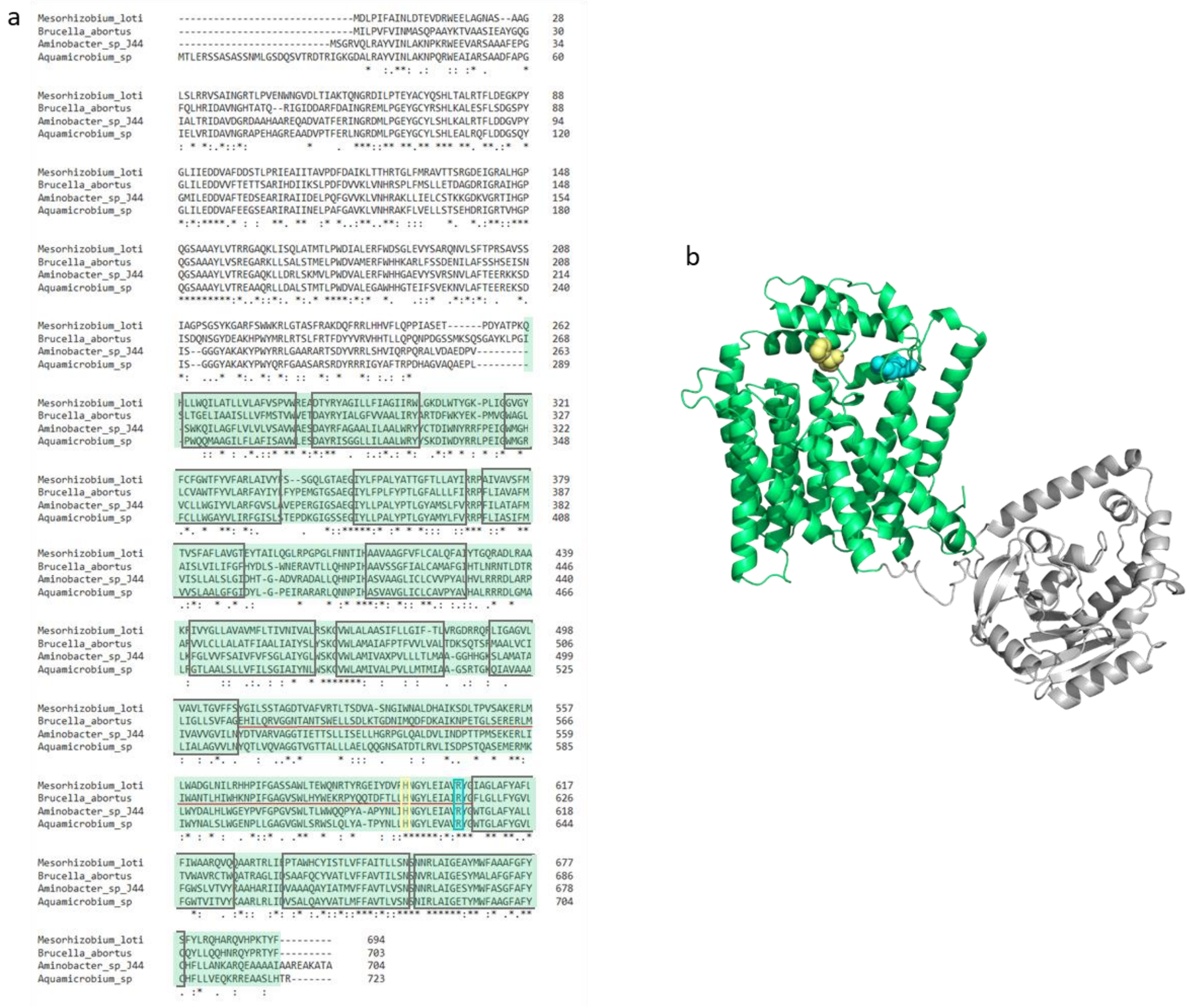


**Figure S6 | Sequence alignment and structure prediction for WadA as the O-antigen ligase in *B. abortus*. a,** Sequence alignment of WadA of *B. abortus*, and WadA homologs of Rhizobiales . Homologs were found using Delta-Blast (Boratyn *et al.*, 2012) and the sequence alignment was performed using Clustal Omega (1.2.4) (Goujon *et al.*, 2010; Sievers *et al.*, 2011). An asterisk (*) indicates a fully conserved amino acid, a colon (:) indicates amino acids with strongly similar properties, and a period (.) indicates amino acids with weakly yet similar properties at the position. The boxes represent transmembrane helixes predicted by DeepTMHMM (v1.0.8) (Hallgren *et al.*, 2022) and the red line is the main periplasmic loop of WadA identified in.*B. abortus.* Arg-614 (R614, cyan) was mutated in this study. Highly conserved His (H605, yellow) was shown to mediate O-antigen ligase activity with a distant homolog (Ashraf *et al.*, 2022). From amino acid 1 to 266 is the glycosyl transferase domain and in green from the amino acid 267 to 703 of *B. abortus* is the predicted O-antigen ligase domain. **b,** Three dimensional model generated by Alpha-fold (Jumper *et al.*, 2021; Varadi *et al.*, 2022) and displayed with PyMol (Schrodinger, 2015). WadA comprises both the glycosyl-transferase (in grey) and the O-antigen ligase domain (in green) which presents 12 transmembrane (TM) segments (grey boxes in the alignment) and 3 periplasmic loops. In cyan, R614 is found in the main periplasmic loop (red line in the alignment), facing the H605 also found in the periplasmic loop.

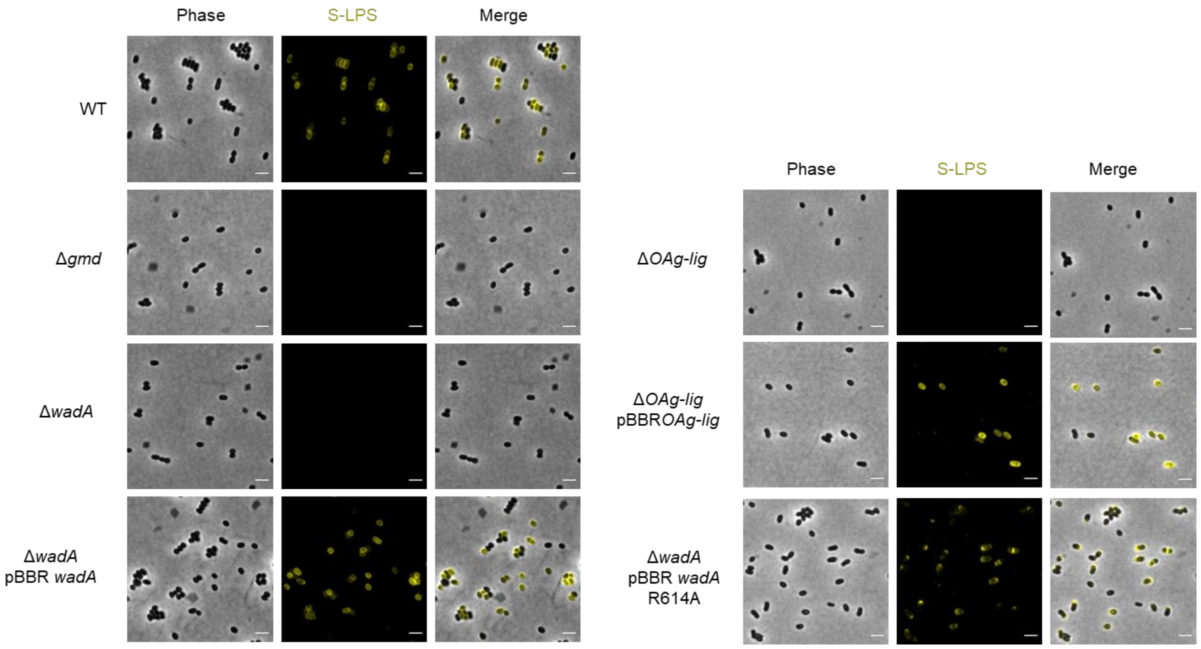


**Figure S7 | S-LPS labelling of *B. melitensis* 16M strains.** *B. melitensis* strains were labelled with primary mAb targeting the O-antigen (A76/12G12) and a goat anti-mouse antibody coupled to Alexa fluor 514 to detect S-LPS. *Δgmd* was used as a negative control. Scale bar is 2µm.
